# STRetch: detecting and discovering pathogenic short tandem repeat expansions

**DOI:** 10.1101/159228

**Authors:** Harriet Dashnow, Monkol Lek, Belinda Phipson, Andreas Halman, Simon Sadedin, Andrew Lonsdale, Mark Davis, Phillipa Lamont, Joshua S. Clayton, Nigel G. Laing, Daniel G. MacArthur, Alicia Oshlack

## Abstract

Short tandem repeat (STR) expansions have been identified as the causal DNA mutation in dozens of Mendelian diseases. Historically, pathogenic STR expansions could only be detected by single locus techniques, such as PCR and electrophoresis. The ability to use short read sequencing data to screen for STR expansions has the potential to reduce both the time and cost to reaching diagnosis and enable the discovery of new causal STR loci. Most existing tools detect STR variation within the read length, and so are unable to detect the majority of pathogenic expansions. Those tools that can detect large expansions are limited to a set of known disease loci and as yet no new disease causing STR expansions have been identified with high-throughput sequencing technologies.

Here we address this by presenting STRetch, a new genome-wide method to detect STR expansions at all loci across the human genome. We demonstrate the use of STRetch for detecting pathogenic STR expansions in short-read whole genome sequencing data with a very low false discovery rate. We further demonstrate the application of STRetch to solve cases of patients with undiagnosed disease and apply STRetch to the analysis of 97 whole genomes to reveal variation at STR loci. STRetch assesses expansions at all STR loci in the genome and allows screening for novel disease-causing STRs.

STRetch is open source software, available from github.com/Oshlack/STRetch.

## Background

Short tandem repeats (STRs), also known as microsatellites, are a set of short (1-6bp) DNA sequences repeated consecutively. Approximately 3% of the human genome consists of STRs [1]. These loci are prone to frequent mutations and high polymorphism, with mutation rates 10 to 100,000 times higher than average rates throughout the genome [2]. Dozens of neurological and developmental disorders have been attributed to STR expansions [3]. STRs have also been associated with a range of functions such as DNA replication and repair, chromatin organization, and regulation of gene expression [2, 4, 5].

STR expansions have been identified as the causal DNA mutation in almost 30 Mendelian human diseases [6]. Many of these conditions affect the nervous system, including Huntington’s disease, spinocerebellar ataxias, spinal-bulbar muscular atrophy, Friedreich’s ataxia, fragile X syndrome, and polyalanine disorders [7]. Most tandem repeat expansion disorders show dominant inheritance, with disease mechanisms varying from expansion of a peptide repeat and subsequent disruption of protein function or stability, to aberrant regulation of gene expression [8].

STR expansion diseases typically show genetic anticipation, characterized by greater severity and earlier age of onset as the tandem repeat expands through the generations [9]. In many STR diseases, the probability that a given individual is affected increases with the repeat length. In some cases severity also depends on the gender of the parent who transmitted the repeat expansion [10]. The number and position of imperfect repeat units also influences the stability of the allele through generations [9]. Together, these features can be used to identify patients with a disease of unknown genetic basis that might be caused by an STR expansion.

Historically, STRs have been genotyped using polymerase chain reaction (PCR) and gel electrophoresis. In such cases, PCR is performed using primers complementary to unique sequences flanking the STR. The PCR product is then run on a capillary electrophoresis gel to determine its size. Although this method has been scaled to handle dozens of samples, it is still labour-intensive and costly. Each new STR locus to be genotyped requires the design and testing of a new set of PCR primers, along with control samples.

A number of diseases are known to be caused by any one of multiple variants, including STR expansions, single nucleotide variants (SNVs) or short indels. For example, there are more than ten STR loci in as many genes that are known to cause ataxia [11], as well as SNVs and indels in dozens of genes [12]. For such diseases, this can mean hundreds of dollars spent per STR locus, plus additional costs for SNV and short indel testing. For such conditions there is a clear need for a single genomic test that can detect all relevant disease variants including SNVs, indels and STRs.

The ability to genotype STRs directly from next-generation sequencing (NGS) data has the potential to reduce both the time and cost to reaching diagnosis and to discover new causal STR loci. It is becoming increasingly common to sequence the genomes or exomes of patients with undiagnosed genetic disorders. Currently, the analysis of this data is focused on SNVs and short indels, and while NGS has identified hundreds of new disease-causing genes, to our knowledge no new pathogenic STR expansions have been discovered. STRs are generally only investigated in an ad-hoc manner at known loci if they are a common cause of the clinical phenotype. The ability to screen for STR expansions in next-generation sequencing data gives the potential to perform disease variant discovery in those patients for which no known pathogenic variants are found.

The vast majority of current STR genotyping tools for short-read sequencing data (most notably LobSTR [13], HipSTR [14] and RepeatSeq [15]) are designed to look at normal population variation by looking for insertions and deletions within reads that completely span the STR. These tools are limited to genotyping alleles that are less than the read length, and require sufficient unique flanking sequence to allow them to be mapped correctly. However, for most STR loci causing Mendelian disease in humans, pathogenic alleles typically exceed 100 bp, with pathogenic alleles at some loci in the range 1,000-10,000 bp [16], far exceeding the size cut-off for detection using these algorithms.

One STR genotyper, STRViper [17], which is also designed to detect population variation, has the potential to detect alleles exceeding the read length by looking for shifts in the distribution of insert size from paired reads. This method requires that the insert size distribution has a relatively low standard deviation and is limited to repeats smaller than the insert size with enough flanking sequence to map both pairs to the reference genome. The mean insert size can be as low as 300-400 bp, meaning that for many large pathogenic expansions, there may be very few or no spanning read-pairs. Such methods would therefore have limited utility for the detection of allele sizes expected for pathogenic expansions. Another tool, ExpansionHunter [18], uses read-pair information and recovery of mis-mapped reads to estimate the length of STRs. For known pathogenic sites ExpansionHunter can be used to determine if the length of the STR is in the pathogenic range. This tool was originally developed for the FTDALS1 repeat and only works on a specific set of pre-defined loci, and is therefore not a genome-wide method. Similarly, exSTRa [19] detects expansions in a set of 21 pre-defined loci and requires a set of matched control samples in order to define the statistical probability of an expansion. In contrast to ExpansionHunter, the exSTRa method does not attempt to estimate allele lengths.

While long-read sequencing technologies can potentially sequence through larger repeat loci [20], they are currently far too expensive for clinical use. High error rates and low throughput also make these technologies less suitable for genotyping SNVs and short indels and are thus a poor alternative to short-read sequencing in a clinical setting. Clearly, there is still a great need to be able to detect STR expansions from short read data.

Here we present STRetch; a new method to detect rare expansions at every STR locus in the genome and estimate their approximate size directly from short read sequencing. We show that STRetch can detect pathogenic STR expansions in short-read whole genome sequencing data and can detect expansions at STR loci not known to be pathogenic. We also demonstrate the application of STRetch to solve cases of patients with undiagnosed disease, in which STR expansions are a likely cause.

STRetch is open source software, available from github.com/Oshlack/STRetch.

## Results

### The STRetch method

The STRetch method has been designed to identify expanded STRs from short-read sequencing data and give approximate sizes for these alleles. Briefly, the idea behind STRetch is to first construct a set of additional sequences comprising all possible STR repeat units in the range 1-6 bp. These are then added to the reference genome as “STR decoy chromosomes”. By mapping to this modified genome, STRetch identifies reads that originated from large STR expansions, containing mostly STR sequence, that now preferentially map to the STR decoys. These reads are then allocated back to the genomic STR locus using read-pair information, and the locus is assessed for an expansion using a statistical test based on coverage of the STR. A summary of the STRetch method is presented in Figure 1, with further specifics detailed below.

**Figure 1:**
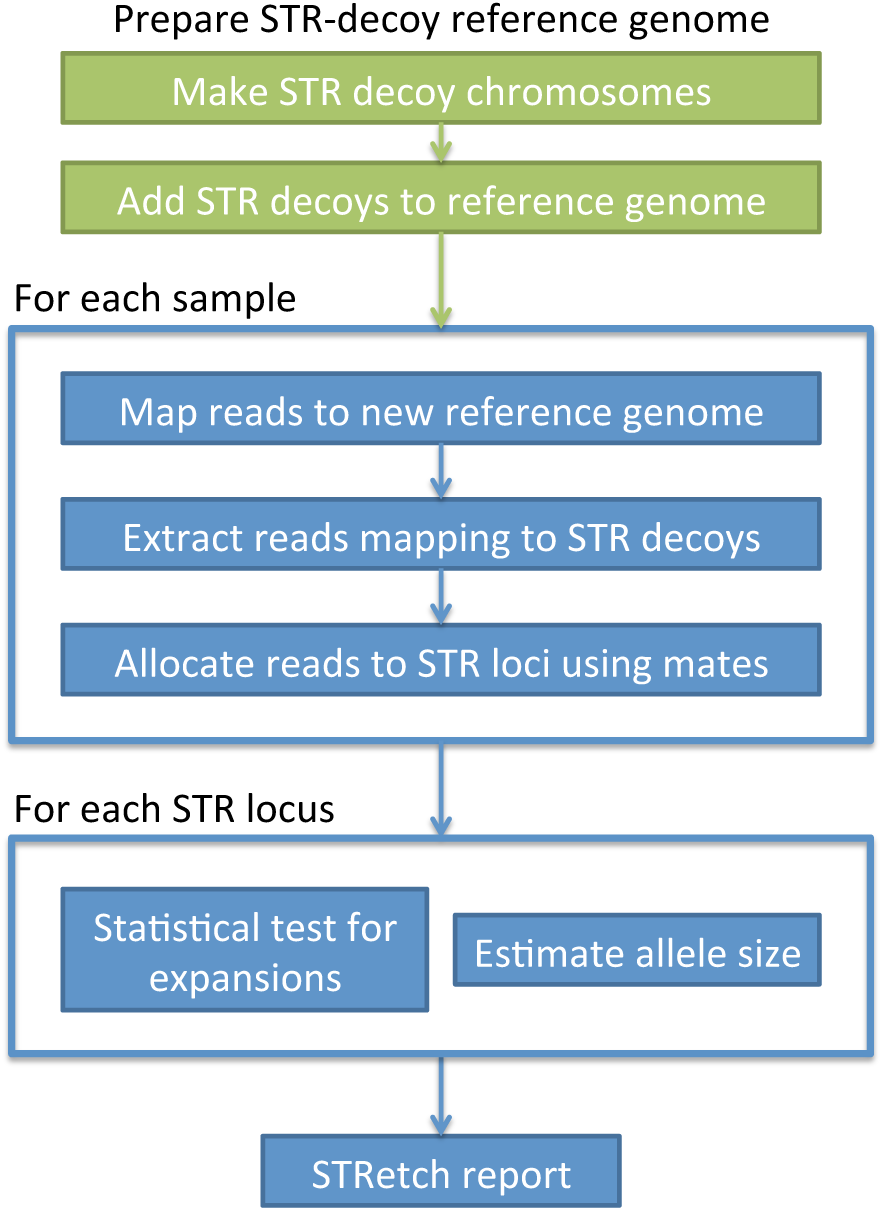
Summary of the STRetch pipeline. The STR decoy reference genome is provided for the user for mapping of each of their test samples. The pipeline will allocate reads to STR loci and perform statistical testing. The resulting report consists of a table with annotation and test results for each locus in each sample.

## STR decoy chromosomes: generating an STR-aware reference genome

A key feature of STRetch is the generation of STR decoy chromosomes to produce a custom STR-aware reference genome. Most aligners have difficulty accurately mapping reads containing long STRs. For example, although the BWA-MEM algorithm has superior performance for mapping reads containing STRs [21], reads containing long STRs sometimes map to other STR loci with the same repeat unit, or completely fail to map [18]. The systematic mis-mapping of STR reads is unsurprising considering that BWA-MEM is optimized to find the longest exact match [22]. For a read made up primarily of STR sequence, the best match is likely to be the longest STR locus in the reference genome with the same repeat unit.

STRetch takes the issue of systematically mis-mapped, or unmapped STR reads and uses this as a way to identify reads that contain long STR sequence. To achieve this, we introduce the concept of STR decoy chromosomes. These are sequences that consist of 2000 bp of pure STR repeat units that can be added to any reference genome as additional chromosomes. STR decoy chromosomes for all possible STR repeat units in the range 1-6 bp are generated and filtered for redundancy, resulting in 501 new chromosomes that are added to the reference genome (STRetch provides hg19 with STR decoys, see Methods). While reads with STR lengths similar to the allele length in the reference genome will map to their original locus, reads containing large STR expansions will preferentially align to the STR decoy chromosomes. These reads are then further examined for evidence of a pathogenic expansion.

## Mapping to STR decoys to identify reads containing STRs

Once the new STR decoy reference genome is created, the first step maps reads against the new reference genome using BWA-MEM. If the data has already been mapped, STRetch can optionally extract and re-map a subset of reads likely to contain STRs. Extracted reads are those that aligned to known STR loci (defined using Tandem Repeats Finder (TRF) [23]), as well as any unmapped reads (see Methods). Any reads mapping to the STR decoy chromosomes are inferred to have originated from an STR. Typically ∼0.01% of reads map to the STR decoy chromosomes in a PCR-free whole genome.

## Determining the origin of STR reads

Next, the reads that map to the STR decoys are assigned to genomic STR positions. STRetch uses the mapping position of the read at the other end of the DNA fragment (the mate read) to infer which STR locus each read originates from. Known STR loci are obtained from a TRF annotation of the reference genome. For a given read, if the mate maps within 500 bp of a known STR locus with the same repeat unit, then the read is assigned to that locus (or the closest matching locus if multiple loci are present). Only 0.93% of STR loci are within 500 bp of another STR locus with the same repeat unit (Supplementary Figure 1). This distance accounts for the fragment length of the majority of reads (Supplementary Figure 2).

After all possible reads are assigned, there may be a difference between the number of reads mapping to a given STR decoy chromosome and the number of reads assigned to all STR loci with that same repeat unit. Unassigned STR reads can occur for a variety of reasons; for example, if their mate also maps to the STR decoy chromosome, if their mate is unmapped, or if their mate does not map in close proximity to a known locus. The number of unassigned reads will increase in samples with very large STR expansions because more read-pairs will originate purely from the STR. This may result in STRetch underestimating the size of very large alleles, however such loci will still be reported as significant as they are still assigned substantially more reads compared with control samples.

## Detecting outlier STR loci

STRetch next uses a statistical test to identify loci where an individual has an unusually large STR. Specifically, STRetch compares the number of STR decoy reads assigned to each locus for a test sample with STR reads from a set of control samples. At each locus the reads are normalized by dividing by the average coverage of the sample. The set of control samples provide a median and variance of counts for each locus. A statistically robust z-score (“outlier score”) is then used to test if the log-normalized number of reads in the test sample is an outlier compared to controls (see Methods). The final result is a multiple-testing-adjusted p-value describing the significance of an expansion at each locus relative to control samples. Every locus for which reads have been assigned is given a p-value.

A variety of control samples can be used in the statistical testing. Firstly, STRetch can be run on a set of controls and the median and variance of the coverage parameters for each locus estimated. These control parameters are then used in testing for significant expansions in disease samples. This approach is ideal for researchers who have access to large sets of sequenced controls. Secondly, a set of samples that are all being tested can be used as controls for each other in a similar way to the above. The assumption here is that only a small proportion of samples (<50%) will have the same expanded locus (we refer to this approach as internal controls). Thirdly, STRetch supplies median and variance parameters estimated from a reference set of PCR-free whole genomes (see “STRetch reveals STR expansions in 97 whole genomes”). This third approach is useful for researchers who only have a limited number of samples to test. The advantage of the first and second options is that the sequencing is usually run at the same centre with the same library preparation protocols. However, the data sets may be smaller and therefore provide less robust statistical measures. In most cases, option three (using the supplied parameters from PCR-free whole genomes) is the easiest and most accurate when using relatively small data sets (see Table 1).

**Table 1:**
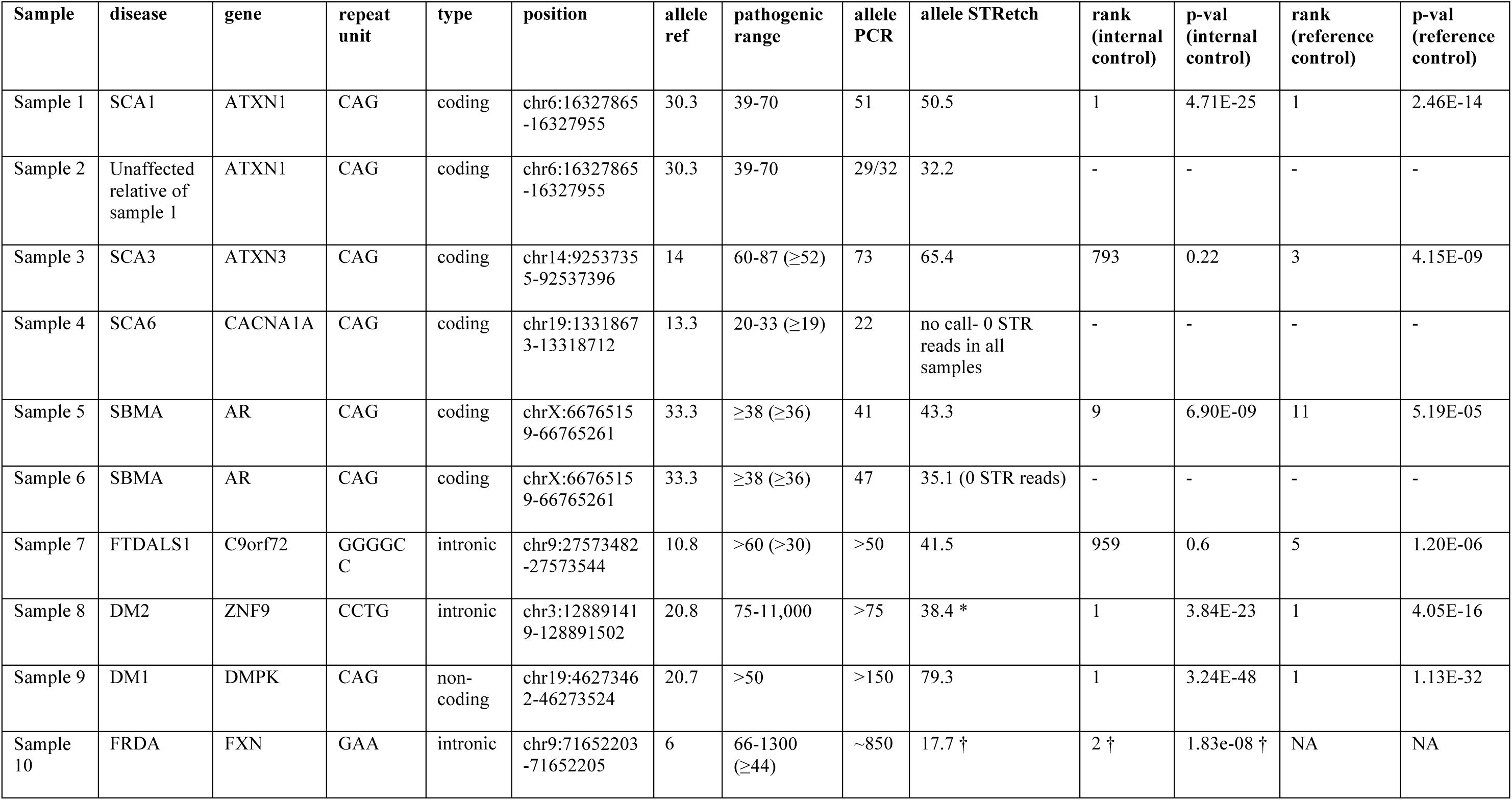
Summary of the 10 individuals with known STR alleles. Allele refers to the number of repeat units. Rank refers to the position of the locus when all the STR loci are ordered based on p-value. Pathogenic range is as reported in GeneReviews [24]. Where there is dispute on the pathogenic range or incomplete penetrance, the alternate range is given in parentheses. * The DM2 locus is a complex repeat: (TG)n(TCTG)n(CCTG)n, where only expansion of the CCTG repeat causes DM2, however all repeat units are polymorphic. STRetch estimates CCTG expansions directly, while the PCR measures the entire complex locus. † Results after manual addition of the FRDA STR to the reference data.

## Estimating the size of STR alleles

When scanning the genome for sites of significant expansions the statistical test is the most important screen. However, for known disease loci the literature has focused on the length of the variant that is associated with pathogenicity. Therefore, STRetch also makes estimates of allele length. STRetch works on the assumption that, for a given locus, the number of reads containing the STR repeat unit is proportional to the length of the repeat in the genome being sequenced. This is because increasing the length of the STR allele increases the likelihood that reads from that locus will be sampled. Hence, STRetch estimates the size of any detected expansion using the normalized read counts allocated to that STR locus. Using simulation, we indeed found that the allocated read counts are linearly related to the length. Specifically, we simulated reads from 100 individuals with the genotype 16xCAG/NxCAG at the SCA8 locus, where N was randomly selected in the range 0 to 500 (see Methods). Our simulated data exhibits a linear relationship between allele size and the number of reads mapping to the STR decoy chromosome (Supplementary Figure 3). We use the slope and offsets from these simulated expansions in estimating the allele size from the normalized coverage at a locus.

## Output files

On completion, STRetch generates a tab-delimited output file for each sample that contains all STR loci for which STR decoy reads were detected. Further information includes p-values for statistical significance of an expansion, details of the STR locus (position, repeat unit, size in reference), robust outlier z-score, locus read count and the allele length estimate. By default, this file is sorted such that the most significant expansions are ranked at the top.

### STRetch is able to recover true pathogenic expansions

In order to test STRetch, we generated PCR-free whole genome sequencing on ten individuals: nine with known pathogenic STR expansions and one unaffected family member. Samples were sequenced to a mean coverage of 41.74x (range 38.35-49.57x), then processed using the Broad GATK pipeline (mapped to hg38 with BWA-MEM, then processed using the GATK best practices).

For analysis with STRetch, we first extracted reads overlapping all known STR loci annotated by Tandem Repeats Finder (see Methods) and then processed these reads through the STRetch pipeline, using the hg19 reference genome. The STRetch statistics were calculated twice; first using only these ten samples as controls for each other (“internal control”) and then using the 97 whole-genome sequencing (WGS) samples described below as controls (“reference control”). We also ran LobSTR/HipSTR to estimate the size of the short allele in each case. On average STRetch reported 18 significant STR expansions per sample (range 4-33).

For six of the ten samples we had information about the disease and the estimated allele size by PCR. For the other four samples (Sample 7, Sample 8, Sample 9 and Sample 10) we were initially completely blinded to all patient information, including phenotype. Disease and allele size estimates were only revealed to us after we had correctly identified the causal STR expansion in each case. Table 1 summarizes the results of this analysis, while Figure 2 and Supplementary Table 1 show allele size estimates for these samples using STRetch, LobSTR, HipSTR and ExpansionHunter.

**Figure 2:**
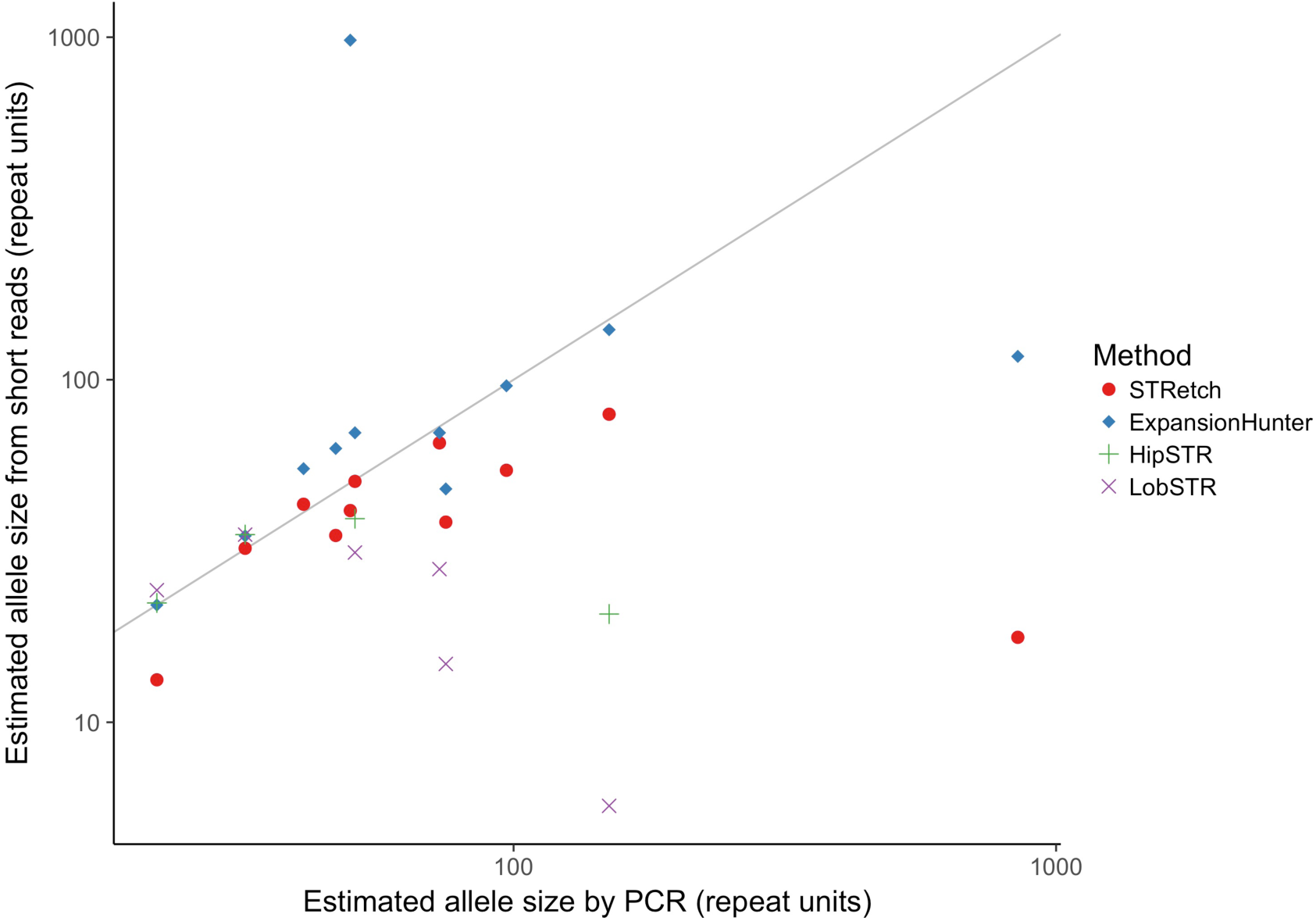
Relationship between allele sizes estimated by PCR and those called by STRetch, ExpansionHunter, HipSTR and LobSTR for the true positive samples. The raw data is available in Supplementary Table 1.

For the six samples with known information, STRetch correctly identified three true positive expansions. Furthermore, STRetch correctly failed to detect an expansion in a pathogenic locus in the true negative (Sample 2). STRetch failed to identify the causal locus in two cases (Sample 4 and Sample 6). For Sample 4 the PCR allele length is only 26 bp larger than the reference, making the entire allele 66 bp, which is well within the read length of 150 bp. STRetch only detects STR expansions that are sufficiently large so that the repeat maps to the STR decoys instead of the reference locus. Indeed, for Sample 4 we see three reads mapped to the genomic locus with a 27 bp insertion, and no reads from this locus mapping to the STR decoy. This allele can be detected by tools that look for indels within the read, and indeed both LobSTR and HipSTR are able to correctly call this expansion (Supplementary Table 1). Sample 6 was found to have lower coverage over the STR region compared with other samples. However, some evidence of the repeat was observed, such as reads ending in the STR with more repeat units than the reference. As such, this expansion may be detected in future iterations of the software.

For the blinded samples (Samples 7, 8 and 9) we were able to correctly determine the causal STR locus simply by ranking variants by their p-values and then looking for any known pathogenic STR loci with a significant p-value (Table 1).

For Sample 10 we were initially unable to identify a significant expansion in a known pathogenic gene. After the variant was revealed to be a large GAA expansion at the FRDA locus we investigated further and discovered that although the reference genome has 6xGAA at this position, the STR is missing from the TRF genome annotation. TRF failed to annotate this STR due to its relatively small size. After manually adding this locus to the genome annotation and rerunning the analysis on the ten samples in Table 1, STRetch was able to correctly detect a significant expansion at this locus. Note that this locus was not annotated in the control samples so only an “internal control” analysis was done.

Most of the true positives that were detected had significant expansions when using both the 97 reference controls and the ten internal controls. However, there were two samples (Samples 3 and 7) that were only significant when using the much larger set of reference controls. Therefore, for this size dataset the use of reference controls generally provides more power.

We compared the PCR sized expansion in these ten samples with the STRetch length estimates and found that STRetch has about a ∼20% standard error. We also found that for very long alleles, STRetch substantially underestimates the allele size. One likely explanation is that the current implementation of STRetch is limited by the insert size of the sequencing data. Some alleles will be so large (e.g. DM1 in Sample 9 is >450 bp) that there will be read-pairs where both are completely contained within the STR. In such cases, both pairs will map to the STR decoy and will not be assigned to an STR locus, leading to underestimation of the read length (Supplementary Figure 4). In comparison, ExpansionHunter more closely estimated alleles in most cases, with a tendency to overestimate allele sizes (Supplementary Table 1). As expected, both LobSTR and HipSTR dramatically underestimated large alleles, or did not make a call in many cases.

STRViper was also run on the true positive samples and reported no significant expansions across all STR loci annotated with TRF. This is likely due to the large standard deviation of insert size for these samples (130-150 bp), which is typical for the current standard Illumina protocol. STRViper requires smaller insert sizes of approximately 30 bp or less.

We next assessed the sensitivity, specificity and false discovery rate (FDR) of STRetch on these samples. We considered STRetch calls in 18 samples for which we had DNA available in our lab (the ten test samples described above, and eight samples from the 97 controls described below) on a set of 22 pathogenic loci (Supplementary Table 2). This gave a test set of 396 loci. We assumed that any significant STRetch calls (p<0.05 after multiple testing correction), beyond the true positive loci described above, are false positives, and the two loci that STRetch failed to detect above are false negatives. Any loci where STRetch does not make a significant call are assumed to be true negatives. 64 of these true negatives were confirmed by PCR from standard diagnostic testing. Using these assumptions, our measured sensitivity was 0.778, specificity was 0.974, and the false discovery rate (FDR) was 0.025. We also ran ExpansionHunter on these samples and found that three of the STRetch calls we had assumed to be false positives were also supported as expanded by ExpansionHunter, although not in the pathogenic range. Two of these (the SCA3 loci) were also confirmed as expanded in the non-pathogenic range by diagnostic PCR. Using these updated values, sensitivity was 0.833, specificity was 0.974 and FDR was 0.018. Six of our ten false positive calls were in the FTDALS locus and another two in the SCA3 locus (although these were expanded, just not-pathogenic). This indicates that some loci may be harder to correctly identify than others. Indeed, the FTDALS1 locus is known to be difficult to analyze due to homology with other loci [18]. Overall, STRetch achieves false discovery rates well below our nominal value of 0.05.

### STRetch reveals STR expansions in 97 whole genomes

We performed PCR-free whole genome sequencing on a set of 97 individuals, most of whom were being investigated for the cause of their Mendelian disease, or were immediate family members of such patients. Many of these cases have inherited neuromuscular disorders, approximately half of which have previously been identified to be caused by a SNV or indel. The remaining cases are still unsolved. The set also includes four individuals with ataxia. In addition to patient samples and relatives, there are seven unaffected samples including NA12878. Given its enrichment for individuals with Mendelian disease, this control set may contain pathogenic STR expansions. However, we did not expect a large proportion of samples to have the same STR expansion so our assumptions for the statistical tests were highly unlikely to be violated.

All samples had previously been mapped with BWA-MEM against hg19. We used STRetch to extract reads from all annotated STR loci, as well as unmapped reads, and re-mapped these against the hg19 STR decoy genome (see Methods). We then proceeded with the rest of the STRetch pipeline. The median and standard deviation were recorded for each locus across all individuals for use as a control set for subsequent analyses. This analysis showed homopolymer loci are the most variable between individuals, showing dramatically higher standard deviations, followed by STRs with 5, 6, 3, 4, and then 2 bp repeat units (Supplementary Figure 5).

To assess the frequency of expansions at known pathogenetic STR loci, we filtered the STRetch results to those significant expansions intersecting with a set of 22 known pathogenic STR loci (Supplementary Table 3). We observed 29 significant STR expansions in seven pathogenic loci (summarised in Table 2). Although these are significantly expanded compared to the rest of the control set, their pathogenicity is uncertain as the allele size estimates are only approximate and are often well below the defined pathogenic range.

**Table 2:**
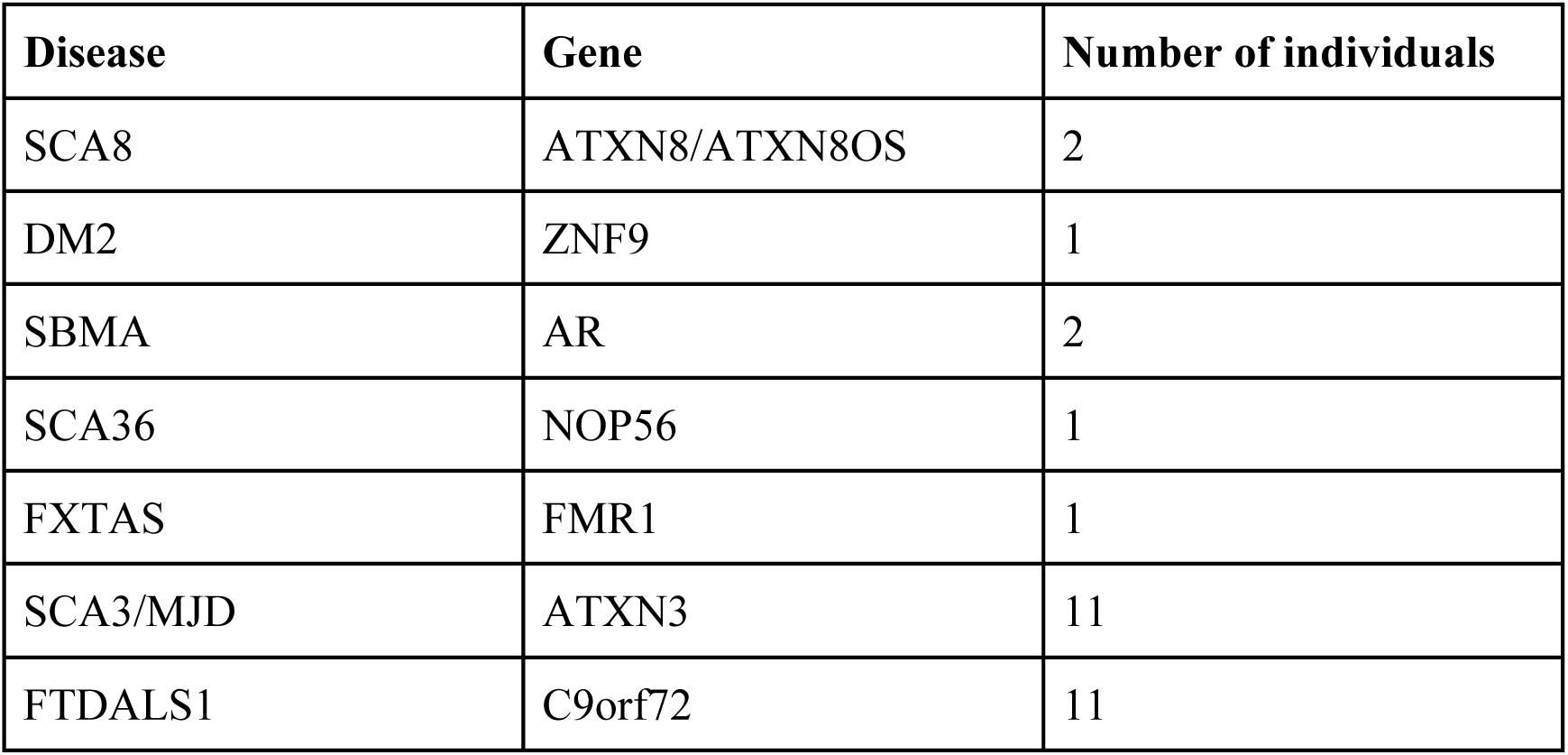
Summary of significant expansions in STR disease loci in 97 WGS samples.

| **Disease** | **Gene** | **Number of individuals** |
| --- | --- | --- |
| SCA8 | ATXN8/ATXN8OS | 2 |
| DM2 | ZNF9 | 1 |
| SBMA | AR | 2 |
| SCA36 | NOP56 | 1 |
| FXTAS | FMR1 | 1 |
| SCA3/MJD | ATXN3 | 11 |
| FTDALS1 | C9orf72 | 11 |

Nonetheless, a number of the STR expansions are potential candidates for follow-up if the sequenced individuals have a relevant phenotype. STRetch detected a large SCA8 expansion in two individuals; one of whom is an ataxia patient (see below). We also detected a DM2 expansion in another individual. All three variants were highly significantly expanded compared to the other control samples (p=4.2e-24, 8.2e-23 and 1.5e-10, respectively), and each was ranked as the most significant for that individual. We have referred these variants back to the originating laboratories to determine if the variants fit the phenotype and can be validated.

STRetch also identified STR expansions in a number of other pathogenic loci in these samples, however many of these are unlikely to be sufficiently large to cause disease. Two SBMA expansions were on the limit of detection and significance (ranked 109 and 476, p=0.001 and 0.04 respectively). We detected SCA36 and FXTAS expansions, which likely reflect sub-clinical variation at these loci, with size estimates of 13.5xAGGCCC and 34.6xCCG respectively. We detected a surprising number of SCA3/MJD_ATXN3 expansions: 11 samples with estimated allele sizes in the range 30.5x to 74.3xAGC. As ≥60xAGC is considered pathogenic, with ≥52 likely showing incomplete penetrance, many of these may be asymptomatic. However, we have observed STRetch to underestimate allele sizes at this locus so these variants could be larger than predicted. We also detected 11 FTDALS1 expansions in the range 20.9-32.9xCCCCGG, all within the unaffected size range for this locus. Generally affected individuals have greater than 60 repeats. Also, smaller allele sizes are associated with later age of onset, with symptoms appearing as late as 80 years old. Consequently, these results may indicate variation within the normal range, individuals with pre-mutations, individuals who have not yet shown symptoms, or false positives. ExpansionHunter confirmed all the SCA8, DM2 and FXTAS expansions, as well as one SBMA, one SCA3 and three of the FTDALS1 expansions. It failed to genotype the SCA36 locus.

These likely non-pathogenic variants at known pathogenic loci highlight the ability of STRetch to explore population variation, and to better determine the true non-pathogenic range of these known pathogenic loci.

### Genetic diagnosis and validation of an ataxia patient

As noted above, four ataxia patients were included in our 97 WGS cohort. These patients had previously been tested with for most known ataxia STR expansions, including SCA1, SCA2, SCA3, SCA6, SCA12, SCA17, SCA38 and DRPLA (see Controls 1-4 in Supplementary Table 2). The whole genome data had also been examined for causative SNVs and indels in candidate ataxia genes using the GATK Best Practices recommendations [25], and copy number variations using BreakDancer [26] and Genome STRiP [27] without success in diagnosis.

In one of these patients (Control 1) STRetch identified the highly significant expansion of SCA8 (noted above), a known but rare disease locus in ATXN8 with an outlier z-score of 11.11 (p=3.55e-24) and an estimated allele size of 54.4x.

As a result, this SCA8 expansion was validated using a PCR assay, which confirmed a pathogenic CAG expansion with an allele size of 97x. In addition, an affected sibling, not sequenced using WGS, was tested for an expansion at the same locus and was similarly determined to have a pathogenic allele of ∼96x. The likely pathogenic range for this STR is 80 to 250 repeat units, with uncertain pathogenicity in the range of 50 to 70 repeat units. However, this locus shows incomplete penetrance, with cases of unaffected individuals observed at all allele sizes [24]. It is noteworthy that STRetch directly estimates the size of the STR, while PCR assays at this locus also include the size of the adjacent non-pathogenic but highly polymorphic CTA repeat. As such, this estimate from STRetch should be interpreted as 65x when comparing to the PCR result.

We also ran LobSTR and HipSTR on the WGS data of this patient. At the expanded pathogenic locus LobSTR called a homozygous 6xTGC insertion, while HipSTR called a homozygous indel from deletion of one TAC repeat unit upstream of SCA8 and an insertion of seven TGC repeat units, for a net total six repeat unit insertion. Both tools report a reference allele size of +15x, so the total allele size is 21x in both cases. PCR analysis of the proband and sibling indicated that both expansions were heterozygous with a short allele size of ∼29x. However, these sizes may not be directly comparable due to potential variation in the definition of the reference locus size.

### STRetch can detect expansions at novel loci

To our knowledge STRetch is the only method currently available that can use short read sequencing to screen the entire genome for rare STR expansions. In order to demonstrate that STRetch can indeed detect expansions at loci not previously associated with disease we performed validation using orthogonal technologies: PCR and Sanger sequencing, and PacBio long-read sequencing.

Firstly, we used PCR to estimate the size of an STR that STRetch called as highly significant in Sample 5. The locus is a 5-base AAACT repeat in an intron of the *MTHFD2* gene (chr2:74430970-74431055). This was the most significantly expanded locus in this sample with a p-value of 4.81E-21 and a predicted size of 37 repeat units. We designed a PCR assay to genotype the size of the allele in this sample as well as five samples that were not predicted to contain this expansion by STRetch (Samples 1, 2, 6, 8 and 9 from above, Supplementary Figure 6). A standard control sample of unknown genotype (CEPH individual 1347-02) was also tested. Sanger sequencing yielded an allele size of greater than 59 repeat units in Sample 5 (the sequencing quality dropped off beyond this size), significantly larger than the alleles from the control samples (p=2.71E-03, Supplementary Figure 7), as predicted by STRetch. By sizing the PCR product on a gel we estimate the larger allele in this sample to be 62 repeat units. We configured ExpansionHunter to estimate the size of this locus resulting in a genotype of 47/64, much larger than the length of 17.2 repeats in the genome. While this STR is unlikely to be pathogenic, this result highlights the potential to use STRetch to screen for novel expansions that may be related to disease.

To further demonstrate the utility of STRetch to identify novel loci as expanded (relative to controls) we compared our analysis of short read data with a previously published genome analysis which utilized long-read PacBio data [28]. Specifically, we ran STRetch on Illumina short read data from an artificial diploid sample created by combining the two haploid genomes, CH1 and CH13 (two replicates), and called significantly expanded loci compared with our 97 reference controls. We then compared all significant STRetch hits with variants called from long-read PacBio sequencing of the two haploid samples. Of the 18 significant STRetch calls (over two replicates) we were able to confirm 14 from the PacBio data as expanded (Supplementary Table 4). Of the four non-validated calls, two were in the non-standard chromosome chrUn_gl000220, which was not present in the PacBio variant data. The remaining two were both homopolymer A STRs. In addition, one of these is in a region with substantial differences between hg19 and hg38 (the genome on which PacBio analysis was performed) including partial deletion of the STR locus, making comparison difficult. While none of the loci represent likely pathogenic expansions, this analysis demonstrates that truly significant expanded variants can be found in novel loci using STRetch with a low false discovery rate.

## Discussion

STRetch is currently the only genome-wide method to scan for STR expansions using short-read whole genome sequencing data. We have demonstrated the ability of STRetch to detect known pathogenic STR expansions in short-read WGS data. STRetch correctly detected the pathogenic expansion in most of our ten test samples; seven true positives and one true negative were correctly identified, and two expansions were missed. The size of one of the missed expansions was below the detection threshold for STRetch. The method performed well on these samples, with a sensitivity of 0.778, specificity of 0.974, and a false discovery rate of 0.025. We applied STRetch to 97 WGS samples to detect both potentially pathogenic STR expansions and expansions of moderate length in STR disease loci, where the allele size is likely below the threshold for disease. Importantly, this set of analyzed genomes can act as a control set, and statistical parameters for the STR loci are provided to use in testing for expansions with STRetch. Within this cohort STRetch revealed a previously undetected pathogenic STR expansion in SCA8 that was validated by PCR in the proband and an affected sibling.

As expected, we found that STR genotypers such as LobSTR and HipSTR, that are designed to genotype short STR variation, were unable to detect large pathogenic variants in WGS data. These tools instead called a homozygous genotype, corresponding to the size of the non-expanded allele, called a heterozygote with slight variation in the small allele, or failed to make a call. Using these tools in conjunction with STRetch allows the estimation of the short allele in cases where STRetch has detected an expansion, allowing us to obtain a more complete picture of the genotype from short read sequencing data.

ExpansionHunter performed relatively well when estimating allele sizes, although tended to overestimate, where STRetch tended to underestimate. However, a key limitation of ExpansionHunter is that it does not use a statistical basis for detecting significantly expanded loci and is not currently configured to estimate lengths across the genome; each locus of interest must be defined in a separate configuration file. Therefore this cannot be used as a genome-wide scan for novel loci.

The main limitation of STRetch is its tendency to underestimate the allele size of STR expansions, especially for variants larger than the insert size. However, this limitation is mostly relevant for known pathogenic loci where there is already an established relationship between the length of the allele and disease characteristics. In genome-wide scans for rare expansions we believe it is more important to use a statistical test that indicates the probability that an expansion exceeds the normal population variation. Estimating allele length by itself does not provide this information. We have demonstrated the potential application of STRetch to discover a novel significant expansion at an intronic STR locus in *MTHFD2*, and validated it by PCR and Sanger sequencing. In addition, further expansions detected by STRetch were also shown to be expanded using long-read sequencing data.

## Conclusions

Here we have introduced STRetch, a method to test for rare STR expansions from whole genome sequencing data. We have shown that STRetch can detect pathogenic STR expansions relevant to Mendelian disease with a low false discovery rate. Although the emphasis has been on STRs known to cause Mendelian disease, a key advantage of STRetch over other methods is its genome-wide approach. STRetch performs statistical tests for expansions at every STR locus annotated in the reference genome, and so has the potential to be used not only for diagnostics, but also in research to discover new disease-associated STR expansions. We hope the application of STRetch to whole genome sequencing of patient cohorts will enable new discoveries of disease-causing STR expansions.

## Methods

### The STRetch pipeline

The STRetch pipeline is implemented using the Bpipe [29] pipeline framework (v0.9.9.3). This allows for a pipeline combining standard bioinformatics tools with novel scripts, and is compatible with many high-performance computing environments, allowing large-scale parallelization over multiple samples.

To summarize the pipeline and components:

Reads are mapped to the reference genome with STR decoy chromosomes using BWA-MEM [22] (v0.7.12) and SAMtools [30] (v1.3.1). STRetch then counts the number of reads mapping to each STR decoy chromosome using bedtools [31] (v2.26.0). Reads mapping to the STR decoy chromosomes are allocated to an STR locus using paired information (Python v3.5.2 script: identify_locus.py). Median coverage over the whole genome or exome target region is calculated using goleft covmed [32] (v0.1.8), which is later used to normalize the counts. STRetch then predicts the size of the expansion using the number of reads allocated to the locus (Python script: estimateSTR.py).

The STRetch pipeline is freely available under an MIT license from github.com/Oshlack/STRetch.

### Generating STR decoy chromosomes

To produce STR decoy chromosomes STRetch generates a set of all possible STR repeat units in the range 1-6 bp. These are then grouped by those repeat units that are equivalent as a circular permutation of each other or the reverse complement. For each group the first repeat unit lexicographically was taken to represent that group. For example, CAG = AGC = GCA = CTG = TGC = GCT, and the group is represented by AGC. STRetch filters out repeat units that could be represented by multiples of a shorter repeat unit. For example, ATAT would be filtered out as it is already represented by AT. This resulted in 501 unique repeat units. STRetch then uses an “STR decoy chromosome” for each repeat unit, which consists of the repeat unit repeated in tandem for 2000 bp (script: decoy_STR.py). This length was chosen to well exceed the insert size, but is configurable. The additional chromosomes can be added to any reference genome (hg19 with STR decoy chromosomes was used for all analyses).

### Extracting likely STR read-pairs from aligned BAMs

In the case where reads have already been mapped to a reference genome, STRetch provides the option of extracting likely STR read-pairs from the BAM file for analysis, rather than remapping all reads in the sample.

In this case, STRetch defines a region where STR reads are likely to align by taking the Tandem Repeats Finder [23] annotation of the reference genome and expanding the region to include 800 bp of flanking sequence on each side. Reads aligned to this region, and all unmapped reads, are extracted using SAMtools view. These are sorted to place together read-pairs using SAMtools collate and then are extracted in fastq format using bedtools bamtofastq. Unpaired reads are discarded.

### Aligning and allocating reads to STR loci

STRetch uses BWA-MEM to align reads to the custom reference genome. Any read mapping to the STR decoy chromosomes is presumed to have originated from an STR locus.

To determine which STR locus the reads originated from, the mates of the reads mapping to a given STR decoy chromosome are collected. If the mate maps within 500 bp of a known STR locus with the same repeat unit, it is assigned to that locus. If multiple loci fall in this range, the read is assigned to the closest locus. This distance was chosen because the mean insert size of our WGS data was 372-415 bp (Supplementary Figure 2), so we expect the mate to map within 500 bp or less of the STR locus. We found that increasing this value did not increase the number of reads allocated to true positive loci. Another consideration when setting this parameter is the potential to misallocate reads from neighbouring STR loci. To combat this, reads are only allocated to a locus with a matching repeat unit. Although rare, there are instances of two STR loci with the same repeat unit occurring close together. In hg19 0.93% of STR loci are within 500 bp of another STR locus with the same repeat unit, 1.34% within 1000 bp, and 2.04% within 2000 bp (see Supplementary Figure 1). We used 500 bp as the smallest value of this parameter that allows detection of all relevant reads while minimizing inclusion of inappropriate reads. This value can be configured in the pipeline if required.

To correct for library size (total number of aligned reads) the counts for each STR locus are normalized against the median coverage for that sample. Counts are normalized to a median coverage of 100x: log_2_(100*(raw counts + 1)/median sample coverage)). Log_2_ normalized counts are used in subsequent statistical analyses.

### Detecting outliers

To detect individuals with unusually large STRs, STRetch calculates an “outlier score” for each individual at each locus. The outlier score is a z-score calculated using robust estimates, with a positive score indicating the STR is larger than the median.

A robust z-score and p-value is calculated for each locus, *l,* using the log normalized counts. First, the median and variance across all samples for locus, *l,* is estimated using Huber’s M-estimator [33, 34]. This calculation can be performed over all samples in a batch, or a set of control samples (estimates from the set of 97 PCR-free whole genomes are provided with STRetch). We test the null hypothesis that the log counts, *y*_*il*_, at locus *l* for sample *i* is equal to the median log counts at locus *l* for the control samples. The alternative hypothesis is that the median log counts for locus *l* are greater for sample *i* compared to the control samples. Hence for each sample *i* and locus *l* we obtain a robust z-score

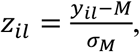

where *M* is the median and *σ*_*M*_ is the square-rooted M-estimator of the variance. One-sided p-values are then obtained from the standard normal distribution and adjusted for multiple testing across the loci using the Benjamini-Hochberg method [35]. A locus is called significant if the adjusted p-value is < 0.05.

### Estimating allele sizes

We reasoned that the size of an expanded allele would be proportional to the number of reads containing STR sequence and hence the number of reads allocated to the STR locus. In order to estimate allele sizes we performed simulations of a single locus at a range of allele sizes. Specifically, reads were simulated from the SCA8 locus in *ATXN8*. One allele was held constant at 16xAGC and then we simulated repeat lengths in the other allele in the range 0-500 repeat units (selected at random from a uniform distribution). Alternate versions of the hg19 reference genome with these alleles were produced using GATK v3.6 FastaAlternateReferenceMaker. Reads were simulated from 10,000 bp either side of the *SCA8* locus using ART MountRainier-2016-06-05 [36]. Reads were 150 bp paired-end, with insert sizes sampled from a normal distribution (mean 500 bp, sd 50 bp) and 30x coverage (proportional coverage sampled from each haplotype). The Illumina error profile was used. Simulation code is available at github.com/hdashnow/STR-pipelines.

A plot of the number of reads mapping to the AGC decoy chromosome against the number of AGC repeat units inserted into the *ATXN8* locus shows a clear linear relationship between these two variables (Supplementary Figure 3).

We can use this information from the simulated data to provide a point estimate of the allele size of any new sample we analyze with STRetch in the following manner. We fit a linear regression between the number of reads mapping to the STR decoy and the size of the allele from the simulated data (both log_2_ transformed), in order to obtain estimates of the intercept and slope parameters, *β*_0_ and *β*,

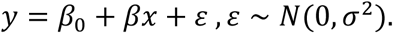

Here y is log_2_(coverage) and x is log_2_(allele size). Given a new data point from a real sample, the log_2_(coverage) for an STR locus of interest, the point estimate of the allele size is thus

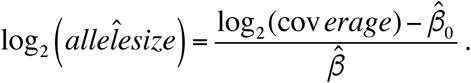

where allele size is the number of base pairs inserted relative to the reference and coverage is the normalized number of reads allocated to the locus.

### Reference data

Reference genome: ucsc.hg19.fasta, with STR decoy chromosomes added as described above.

STR positions in genome annotated bed file: hg19.simpleRepeat.txt.gz. Source: http://hgdownload.cse.ucsc.edu/goldenPath/hg19/database/

Known STR loci are obtained by performing a Tandem Repeats Finder [23] annotation of the reference genome. Pre-computed annotations of many genomes are available from the UCSC Table Browser (https://genome.ucsc.edu/cgi-bin/hgTables) [37]. TRF annotations are converted to bed files annotated with two additional columns: the repeat unit/motif and the number of repeat units in the reference.

### Running other STR genotypers

To estimate the size of the shorter allele, LobSTR and HipSTR were run on BAM files containing the locus of interest and 1000 bp of flanking sequence on either side. We used LobSTR version 4.0.6 with default settings and the LobSTR reference genome and annotation hg19_v3.0.2. HipSTR version 0.4 was used --min-reads 2 and otherwise default settings, with the provided hg19 reference genome and annotation. In some cases, the tools make a different call as to the reference allele in hg19. To make variant calls comparable to STRetch calls we converted genotypes to numbers of repeat units inserted relative to the reference defined by that tool, then applied that to the reference allele given by STRetch. For example, if the STRetch reference allele is 20 repeat units, while in HipSTR it is 10 repeat units, a HipSTR genotype of 10/15 would be reported as 20/25. All imperfect repeat units or other variation was ignored and only the total size of alleles taken.

ExpansionHunter version 2.5.5 was run using the 17 provided STR loci specifications, as well as manually defined loci for STRs in ATXN8, MTHFD2, NOP56 and ZNF9 (see Supplementary Methods). The “original” version of STRViper was downloaded from http://bioinf.scmb.uq.edu.au/STRViper and run using default settings.

### Validation of novel STR expansions

To validate the novel STR expansion in an intron of *MTHFD2* (as predicted by STRetch), PCR was conducted using GoTaq G2 colorless master mix (Promega, USA) with 0.5 µM primers (see Supplementary Table 5) and 10 ng of template DNA per reaction. Cycling was as follows: 95°C for 2 min, 40 cycles of 95°C for 15 s, 68°C for 15 s, and 72°C for 30 s, and a final extension for 5 min at 72°C. Samples were analyzed on a 2% agarose gel stained with ethidium bromide (0.5 µg/mL). Product sizes and STR length were estimated relative to a 100 bp DNA ladder by generating a standard curve (NEB, USA). For sequencing, individual alleles were separated by band-stab PCR [38] (Supplementary Table 5), purified using a QIAquick PCR purification kit (QIAGEN, USA) and sequenced using the Sanger method. The samples tested were Samples 1, 2, 5, 6, 8 and 9 from the true positive samples described above, and one standard control sample 1347-02 (CEPH).

To validate STRetch results with long-read data, STRetch was run on Illumina short read data from an artificial diploid sample created by combining the two haploid genomes, CH1 and CH13. The two replicates were obtained from SRA; https://www.ncbi.nlm.nih.gov/sra, accession numbers ERX1413365 and ERX1413368. We compared the STRetch results to publicly available PacBio variants for these same samples [28]. The authors generated these variants using their SMRT-SV software [24] against the hg38 reference genome. Long structural variants (SVs) >50 bp were obtained from dbVar; https://www.ncbi.nlm.nih.gov/dbvar accession number nstd137 and short SVs from http://eichlerlab.github.io/pacbio_variant_caller/. A STRetch call was considered validated if the PacBio data contained a substantial expansion relative to the reference genome at the same position with the same repeat unit.

## Availability of data and material

The STRetch software is available from https://github.com/Oshlack/STRetch. Installation of the software includes reference data and summary statistics from control samples. These are also available separately from https://github.com/Oshlack/STRetch/wiki. PCR-free whole genome sequencing of the ten test samples is available under BioProject (https://www.ncbi.nlm.nih.gov/bioproject) accession PRJNA419676.

